# Targeted directional kilobase sequence insertion by combining prime editing with recombinases or integrases

**DOI:** 10.1101/2022.05.25.493515

**Authors:** Nathaniel Jillette, Jacqueline Jufen Zhu, Albert Wu Cheng

## Abstract

Targeted insertion of exogenous sequences to genomes is useful for therapeutics and biological research. While CRISPR/Cas technologies have been very efficient in gene knockouts by double-strand breaks (DSBs) followed by indel formation through non-homologous end-joining (NHEJ) repair pathway, the precise introduction of new sequences mainly rely on inefficient homology directed repair (HDR) pathways following Cas9-induced DSBs and are restricted to dividing cells. The recent invention of Prime Editing allows short sequences to be precisely inserted at target sites without DSBs. Here, we combine Prime Editing and sequence-specific recombinases and integrases to insert kilobase sequences directionally at target sites. This technique, called PRIMAS for <u>P</u>rime editing, <u>R</u>ecombinase, Integrase-<u>m</u>ediated <u>A</u>ddition of <u>S</u>equence, will expand our genome editing toolbox for targeted insertion of long sequences up to kilobases and beyond.

## Introduction

CRISPR/Cas technology has provided great power in editing the genome^1–3^. While indel-mediated knockout of genes is efficient with the NHEJ repair pathway following double-stranded breaks (DSBs), precise editing relies on homologous recombination happening at a much lower efficiency and is restricted mostly to dividing cells^4^. Furthermore, the reliance on DSBs means that off-target cutting by Cas9 might induce unwanted perturbations in the genome. Prime editing is a recent advancement in the CRISPR/Cas toolbox using Cas9 nickase to generate single-stranded nick; the nicked strand subsequently acts as primer annealing to an extended prime editing guide RNA (pegRNA) which serves as a template for reverse transcription (RT)^5^. Sequence changes or insertions are encoded in the template region of pegRNA and are incorporated at the target. Prime editing allows addition of sequences without DSBs in non-dividing cells, thus improving the safety and scope of CRISPR/Cas genome editing. Prime editing has been shown to insert short sequences, however, longer sequences up to thousands of bases have not been demonstrated. On the other hand, site-specific recombinases and integrases (e.g., BxB1, PhiC31, Cre, FlpE) can mediate insertion of DNA via specific sequences and have been widely applied in genetic models^6–9^. The insertion of DNA is mediated by short sequences which might be amenable to prime editing-mediated insertion. Here, we describe a hybrid approach, PRIMAS (<u>P</u>rime editing, <u>R</u>ecombinase, Integrase-<u>m</u>ediated <u>A</u>ddition of <u>S</u>equence), using a combination of prime editor and recombinase/integrase to insert kilobases of DNA into target site. Briefly, recombinase/integrase recognition sites are encoded on a pegRNA and inserted to the target site via prime editing. A recombinase/integrase uses the inserted recognition site at the target locus to incorporate donor DNA flanked by cognate recombinase/integrase sites.

## Results

### Targeted directional insertion of donor DNA payload by combining prime editing and dual integrases

We tested PRIMAS strategy at the HEK3 locus for inserting a donor DNA payload containing CAGGS promoter-driven Blasticidin resistance gene-2A-mScarlet fluorescent gene-BGH polyA signal (CAGGS-Blast-2A-mScarlet-pA, 2605bp). Many applications such as protein tagging require insertion of DNA with directional specificity. To achieve that, we tested if we could use two different integrases to flank the payload DNA for use in PRIMAS. We created a donor plasmid with AttB(Bxb1) and AttB(PhiC31) sites^8^ flanking the payload (**FIG. 1A**). We constructed a vector expressing pegRNA with priming site for HEK3 locus and RT template encoding attachment sites for Bxb1 and PhiC31, i.e., AttP(Bxb1) and AttP(PhiC31), respectively (**FIG. 1A**). We transfected HEK293T cells with Prime editor 2^5^ (pCMV-PE2, Cas9(H840A)-M-MLV RT(D200N/L603W/T330P/T306K/W313F), Addgene #132775), BxB1-expressing plasmid^8^ (pCMV-Bx, Addgene #51552), PhiC31-expression plasmid^8^ (pCS-kI, Addgene #51553), pegRNA-expressing plasmid, and the donor DNA vector (**FIG. 1A**). To detect the targeted insertion of the DNA payload, we harvested cells 48 hours after transfection, extracted genomic DNA, and conducted genotyping PCR using primers P1 and P2, which anneal to HEK3 genomic target and payload DNA, respectively. Sample transfected with the full set of plasmids (EXP: experimental sample) show a band indicative of the insertion while control sample (CTL) transfected with a pegRNA without AttP sites is negative for the PCR (**FIG. 1B**). Sequencing of the purified PCR band from the experimental sample confirmed the expected HEK3 genomic sequence, recombined AttR sequence, as well as vector sequence indicative of the precise insertion of the DNA payload (**FIG. 1B**).

**FIG. 1.**
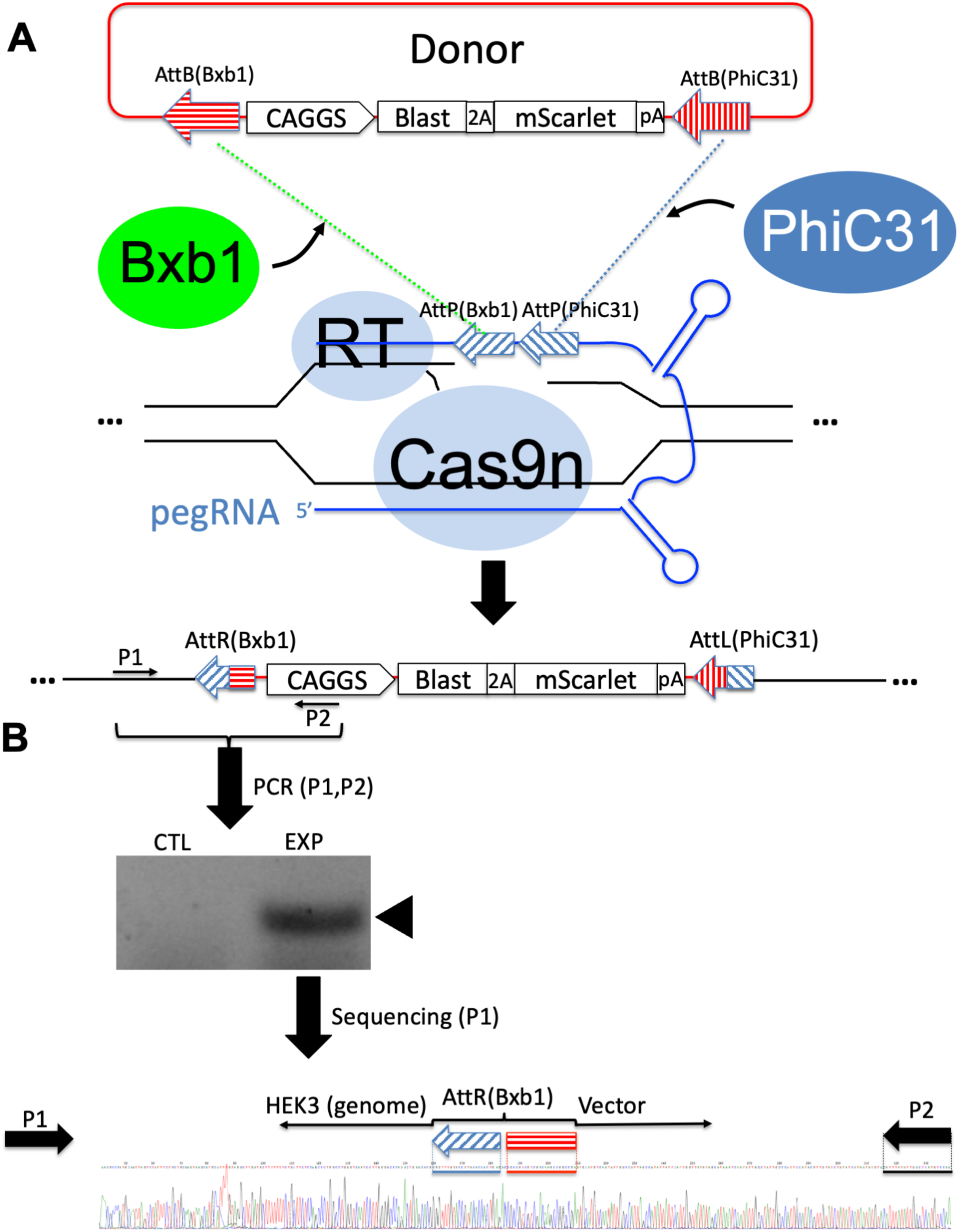
Targeted insertion of donor DNA payload by combining prime editing and dual integrases. **(A)** Templated addition of a Bxb1 AttP site and a PhiC1 AttP site encoded on the prime editing guide RNA (pegRNA) by prime editor 2 (PE2, Cas9n-RT) at a target locus (e.g., HEK3) allows Bxb1- and PhiC1-mediated site-specific recombination with cognate AttB site pairs flanking a payload sequence on a donor vector, resulting in donor payload sequence insertion at the target locus. **(B)** Genotyping PCR using genomic target-specific primer (P1) and vector-specific primer (P2) revealed the presence of targeted insertion product in experimental sample (EXP) absent from control (CTL) sample which had received a pegRNA without the AttP sites. Sequencing trace shows the precise junctional sequences derived from the genomic target and donor vector and the recombined Bxb1 AttR (AttP x AttB) site.

### Targeted directional insertion of donor DNA payload by combining prime editing and BxB1 integrase directed by orthogonal flanking attachment sites

While the use of dual integrases could ensure directionality of payload insertion, the requirement for the delivery of two integrases simultaneously may pose delivery challenge for future therapeutic applications. Bxb1 integrase is known to tolerate dinucleotide mutations at the center of the integration sequence^6^. In addition, the dinucleotide variants are known to be orthogonal. We thus set out to test if we could flank payload DNA with orthogonal Bxb1 site variants to achieve directional insertion with only the Bxb1 integrase. We created a donor DNA vector with payload-flanking orthogonal AttB(gt) and AttB(ga) sites harboring GT and GA central dinucleotide, respectively (**FIG. 2A**), and a vector expressing pegRNA with priming site for HEK3 locus and RT template encoding the corresponding AttP(gt) and AttP(ga) sites (**FIG. 2A**). We transfected HEK293T cells with Prime editor 2 (pCMV-PE2), BxB1-expressing plasmid (pCMV-Bx), pegRNA-expressing plasmid, and the donor DNA vector (**FIG. 2A**). Sample transfected with the full set of plasmids (EXP: experimental sample) showed a genotyping PCR band indicative of the insertion which was negative in control sample (CTL) transfected with a pegRNA without AttP sites (**FIG. 2B**). Sequencing of the positive band from the experimental sample confirmed the expected HEK3 genomic sequence, recombined AttR sequence, as well as vector sequence resulting from the precise insertion of the DNA payload (**FIG. 2B**).

**FIG. 2.**
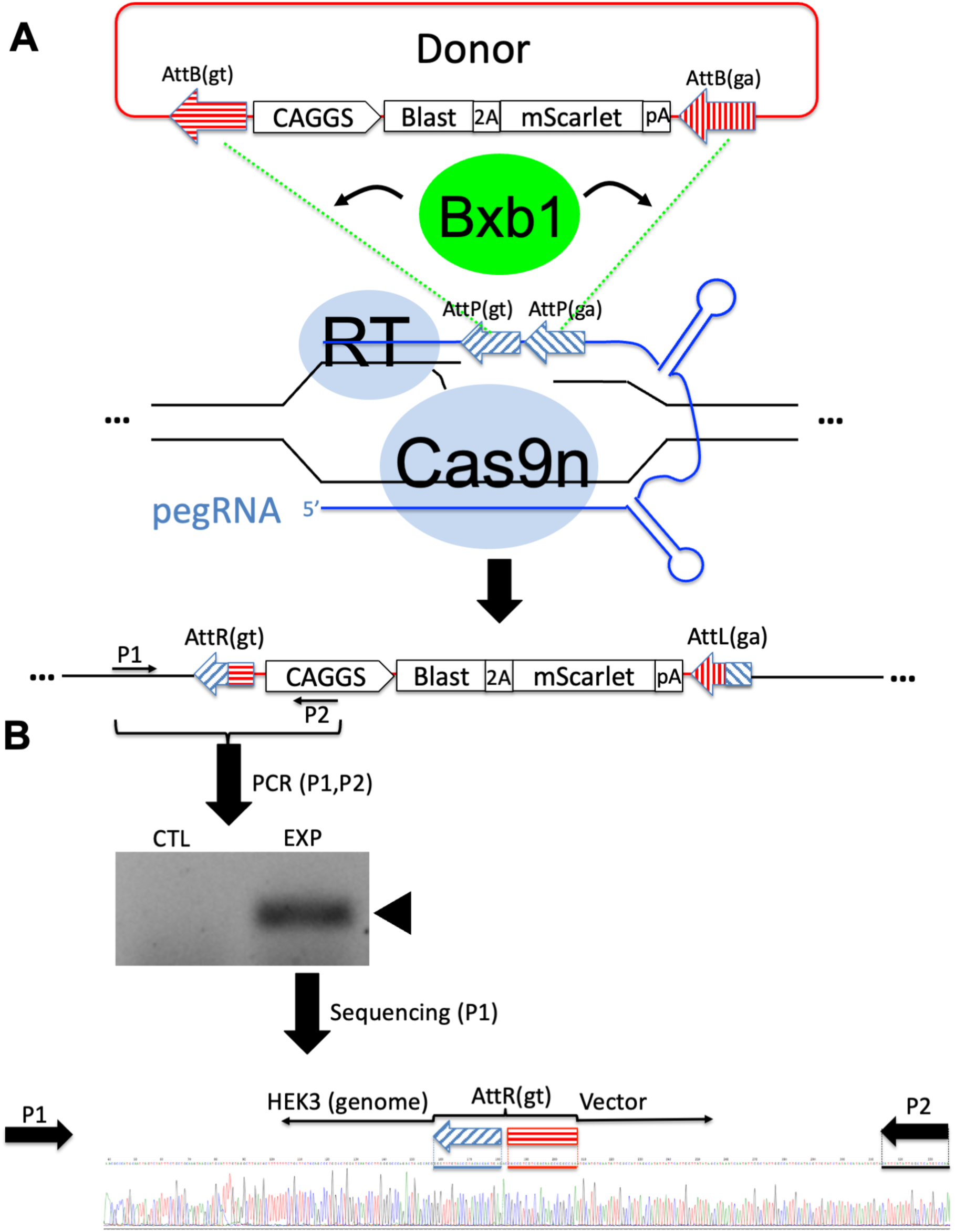
Targeted insertion of donor DNA payload by combining prime editing and BxB1 integrase. **(A)** Templated addition of orthogonal Bxb1 AttP site pairs with GA and GT central dinucleotide encoded on the prime editing guide RNA (pegRNA) by prime editor 2 (PE2, Cas9n-RT) at a target locus (e.g., HEK3) allows Bxb1-mediated site-specific recombination with cognate AttB site pairs flanking a payload sequence on a donor vector, resulting in donor payload sequence insertion at the target locus. **(B)** Genotyping PCR using genomic target-specific primer (P1) and vector-specific primer (P2) revealed the presence of targeted insertion product in experimental sample (EXP) absent from control (CTL) sample which had received a pegRNA without the AttP sites. Sequencing trace shows the correct junctional sequence derived from the genomic target and donor vector as well as the recombined AttR (AttP x AttB) site.

### Targeted insertion of donor DNA payload by combining prime editing and FlpE recombinase

We next tested if recombinases such as FlpE could be used for payload insertion in the context of PRIMAS. We created a donor plasmid with two FRT sites^7^ flanking the Blast-2A-mScarlet payload (**FIG. 3**), and a vector expressing pegRNA with priming site for HEK3 locus, and RT template encoding an FRT site^7^ (**FIG. 3A**). We transfected HEK293T cells with Prime editor 2 (pCMV-PE2), FlpE-expressing plasmid^7^ (pCAGGS-FlpE-puro, Addgene #20733), pegRNA-expressing plasmid, and the donor DNA vector (**FIG. 3A**). Sequencing of the genotyping PCR product from the experimental sample confirm the expected HEK3 genomic sequence, FRT sequence, as well as vector sequence indicative of the precise insertion of the DNA payload (**FIG. 3B**).

**FIG. 3.**
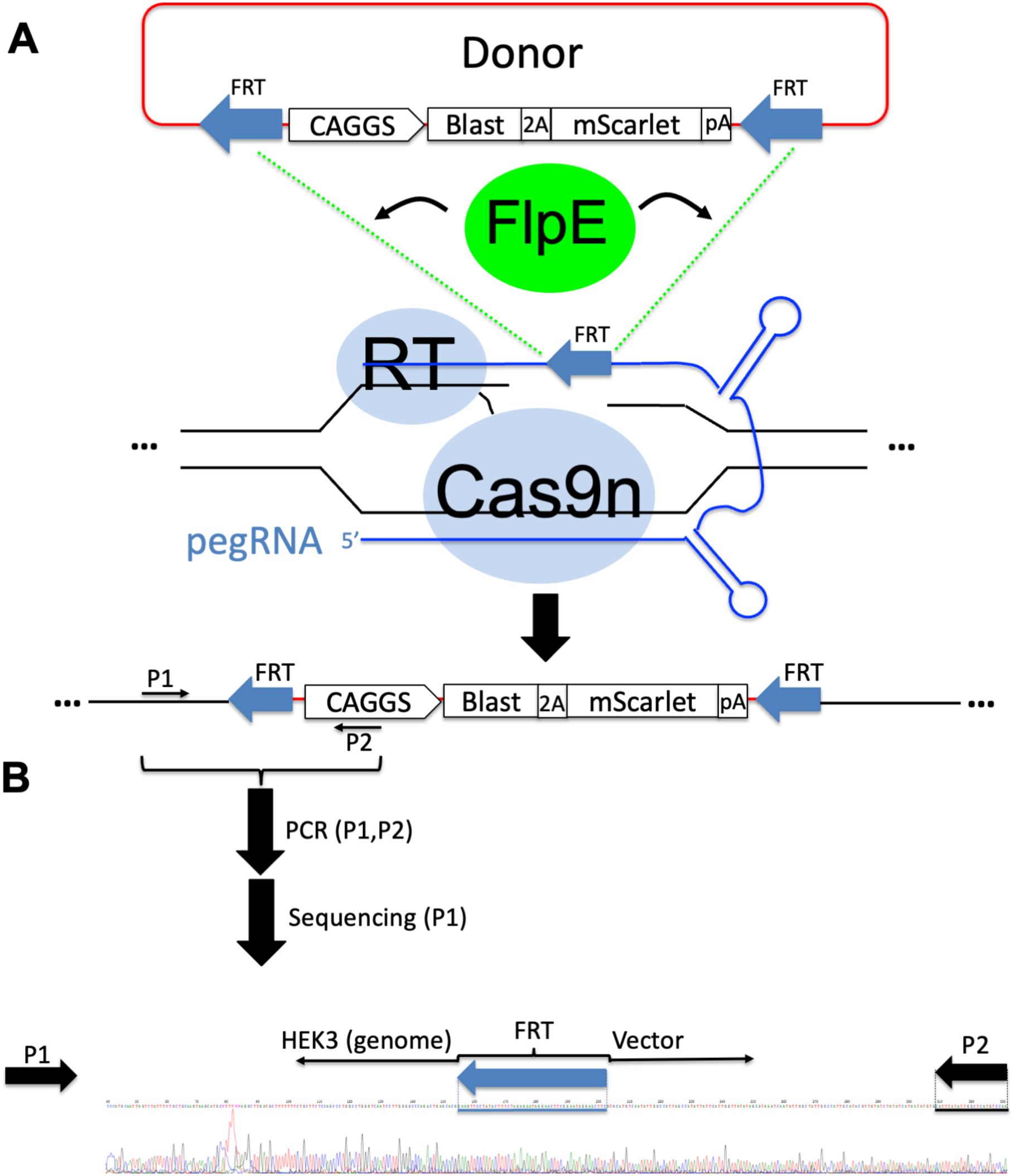
Targeted insertion of donor DNA payload by combining prime editing and FlpE recombinase. **(A)** Templated addition of an FRT site encoded on the prime editing guide RNA (pegRNA) by prime editor 2 (PE2, Cas9n-RT) at a target locus (e.g., HEK3) allows site-specific recombination with FRT site pairs flanking a payload sequence on a donor vector, resulting in donor payload sequence insertion at the target locus. **(B)** Sequencing trace shows the precise junctional sequences derived from the genomic target and donor vector and the reconstituted FRT site.

## Discussion

Targeted insertion of long DNA sequences such as therapeutic transgene at safe harbor or the in-frame tagging of fluorescent proteins are powerful techniques for gene therapy as well as biological research. The reliance of low efficiency homology-directed repair (HDR) methods which additionally requires dividing cells limit their scope of applications. The recent invention of Prime Editing technologies allows efficient insertion of short sequences at defined targets in dividing and non-dividing cells. Here, we demonstrate the PRIMAS approach - the use of prime editing to install recognition sites for recombinases and integrases, and concomitant recombinase- and integrase-mediated directional payload DNA insertion. We believe PRIMAS will be a powerful addition to our genome editing toolbox that allows double-stranded break-independent directional insertion of long sequences up to kilobases and beyond.

## Materials & Methods

### Cloning and sequence listing

pCMV-PE2^5^ (Addgene #132775), pCMV-Bx^8^ (Addgene #51552), pCS-kI^8^ (Addgene #51553) pCAGGS-FlpE-puro^7^ (Addgene #20733) were obtained from Addgene. PegRNA containing integrase/recombinase sites were constructed by a combination of PCR using Ultramers ordered from IDT, and ligation free cloning method TEDA^10^. Sequences of different elements are listed below:

~~~
>AttB(gt) = AttB(Bxb1) wildtype sequence with gt central dinucleotide
TCGGCCGGCTTGTCGACGACGgcggtctcCGTCGTCAGGATCATCCGGGC
>AttB(ga) = AttB(Bxb1-ga) mutant site with ga central dinucleotide
TCGGCCGGCTTGTCGACGACGgcggactcCGTCGTCAGGATCATCCGGGC
>pegRNA-HEK3_BxB1ga-BxB1_attP ggcccagactgagcacgtgagtttAagagctaTGCTGGAAACAGCAtagcaagttTaaataaggctagtccgttatc
aacttgaaaaagtggcaccgagtcggtgctctgccatcaGTCGTGGTTTGTCTGGTCAACCACCgcggactcAGTGG
TGTACGGTACAAACCCCGACgagttggtcgtcgtaccgtaTCGTGGTTTGTCTGGTCAACCACCGCGgtCTCAGTGG
TGTACGGTACAAACCCcgtgctcagtctgttttttt
>AttP(gt) = AttP(Bxb1) wildtype sequence with gt central dinucleotide
TCGTGGTTTGTCTGGTCAACCACCGCGgtCTCAGTGGTGTACGGTACAAACCC
>AttP(ga) = AttP(Bxb1-ga) mutant site with ga central dinucleotide
GTCGTGGTTTGTCTGGTCAACCACCgcggactcAGTGGTGTACGGTACAAACCCCGAC
>pegRNA without AttP sites
ggcccagactgagcacgtgagtttAagagctaTGCTGGAAACAGCAtagcaagttTaaataaggctagtccgttatc
aacttgaaaaagtggcaccgagtcggtgctctgccatcaaagcgtgctcagtctgttttttt
>AttB(PhiC31)
GTGCGGGTGCCAGGGCGTGCCCTTGGGCTCCCCGGGCGCGTACTCC
>pegRNA-HEK3_PhiC1-BxB1_attP ggcccagactgagcacgtgagtttAagagctaTGCTGGAAACAGCAtagcaagttTaaataaggctagtccgttatc
aacttgaaaaagtggcaccgagtcggtgctctgccatcaCCCAGGTCAGAAGCGGTTTTCGGGAGTAGTGCCCCAAC
TGGGGTAACCTTTGAGTTCTCTCAGTTGGGGGCGTAGGGTCGCCGACATGACACAAGGGGTTgagttggtcgtcgta
ccgtaTCGTGGTTTGTCTGGTCAACCACCGCGgtCTCAGTGGTGTACGGTACAAACCCcgtgctcagtctgtttttt
t
>AttP(PhiC31)
CCCAGGTCAGAAGCGGTTTTCGGGAGTAGTGCCCCAACTGGGGTAACCTTTGAGTTCTCTCAGTTGGGGGCGTAGGG
TCGCCGACATGACACAAGGGGTT
>FRT
GAAGTTCCTATTCcGAAGTTCCTATTCtctagaaaGtATAGGAACTTC
>pegRNA-HEK3_FRT
ggcccagactgagcacgtgagtttAagagctaTGCTGGAAACAGCAtagcaagttTaaataaggctagtccgttatc
aacttgaaaaagtggcaccgagtcggtgctctgccatcaGAAGTTCCTATTCcGAAGTTCCTATTCtctagaaaGtA
TAGGAACTTCcgtgctcagtctgttttttt
~~~

### Cell culture and transfection, Genotyping PCR and sequencing

HEK293T cells were cultivated in Dulbecco’s modified Eagle’s medium (DMEM) (Sigma) with 10% fetal bovine serum (FBS)(Lonza), 4% Glutamax (Gibco), 1% Sodium Pyruvate (Gibco) and penicillin-streptomycin (Gibco). Incubator conditions were 37 °C and 5% CO2. Cells were seeded into 24-well plates the day before transfection. Cells were transfected with 600ng total plasmid DNA using Lipofetamine 3000 reagent (Invitrogen). For experiment shown in Fig 1, the plasmid mix included 67ng each of pCMV-PE2, pCMV-Bx, pCS-kI, as well as 200ng of pU6-pegRNA-HEK3_PhiC1-BxB1_attP and 200ng of attB(Bxb1)_Blast-P2A-mScarlet_attB(PhiC31). In the control transfection, a pegRNA inserting CTT triplet^5^ instead of integrase sites was used. For experiment shown in Fig 2, the plasmid mix included 100ng of pCMV-PE2, 100ng of pCMV-Bx, 200ng of pU6-pegRNA-HEK3_BxB1ga-BxB1_attP, and 200ng attB(Bxb1GT)_Blast-P2A-mScarlet_attB(Bxb1GA). For experiment shown in Fig 3, 100ng of pCMV-PE2, 100ng of pCAGGS-Flpe-puro, 200ng of pU6-pegRNA-HEK3_FRT and 200ng of FRT_Blast-P2A-mScarlet-FRT were used. The day after transfection, cells were replenished with fresh media. 48 hours after transfection, cells were harvested and subjected to DNeasy Blood and Tissue Kit (QIAGEN) for genomic DNA extraction. Extracted genomic DNA was genotyped for targeted insertion with primers, P1 (ATGTGGGCTGCCTAGAAAGG) and P2 (TTGGACATGAGCCAATATAAATG) using Phusion DNA polymerase (Thermo Fisher). The positive band from genotyping PCR was subjected to Sanger sequencing.

## Acknowledgements

pCMV-PE2 (Addgene #132775) was a gift from David Liu. pCMV-Bx (Addgene #51552), pCS-kI (Addgene #51553) were gifts from Michele Calos. pCAGGS-FlpE-puro (Addgene #20733) was a gift from Rudolf Jaenisch. This work has been supported by the National Human Genome Research Institute grant R01HG009900 (to A.W.C). Arizona State University startup grant (to A.W.C.) and Jackson Laboratory internal grants (to A.W.C.).

